# WBC-Profiler: an Unsupervised Feature Learning System for Leukocytes Characterization and Classification

**DOI:** 10.1101/508564

**Authors:** Hong Yan, Xuanyu Mao, Yongquan Xia, Zhiyang Li, Chengbin Wang, Rui Xia, Xuejing Xu, Zhiqiang Wang, Xie Zhao, Yan Li, Han Shen, Hang Chang

## Abstract

The characterization and classification of white blood cells (WBC) is critical for the diagnosis of anemia, leukemia and many other hematologic diseases. We developed WBC-Profiler, an unsupervised feature learning system for quantitative analysis of leukocytes. We demonstrate that WBC-Profiler enables automatic extraction of complex signatures from microscopic images without human-intervention and thereafter effective construction of leukocytes profiles, which decouples large scale complex leukocytes characterization from limitations in both human-based feature engineering/optimization and the end-to-end solutions provided by modern deep neural networks, and therefore has the potential to provide new opportunities towards meaningful studies/applications with scientific and/or clinical impact

## Introduction

Hematology tests provide laboratory assessments of blood formation and blood disorders, and play critical roles in the indication, diagnosis and evaluation of many conditions, including infection, inflammation and anemia. Among various hematology tests, the differential count of while blood cells (WBC) provides an important tool in diagnosing and monitoring infection and leukemic disorders, and the ratio of various kinds of leukocytes are commonly used as important markers. For example, the neutrophil to lymphocyte ratio (NLR) is used as a marker of subclinical inflammation, and recent studies suggest that increased NLR is independent predictor of mortality in patients undergoing angiography or cardiac revascularization [1], meanwhile it is also associated with poor prognosis of various cancers [2], such as esophageal cancer [3] or advanced pancreatic cancer [4].

Accurate characterization, detection and classification of while blood cells into several categories, including Monocytes, Lymphocytes, Basophils, Eosinophils, Atypical lymphocytes and Neutrophilic granulocytes, is critical for the ratio (proportion) assessment of leukocytes in blood cell slides. However, such a high-demanding task in clinical laboratory heavily relies on the manual annotation by pathologies, which is not only labor-intensive but also challenging given the shortage of experienced medical experts in many clinical laboratories. To overcome these obstacles, many efforts have been made towards the automatic blood cell classification. Among which, early development of commercial systems in 1970s failed to revolutionize the field due to the high price and low accuracy [5] of the products, and Leuko, another commercial leukemia diagnosis system, was later on developed to reveal an improved accuracy on cell classification based on naive Bayes classifiers. Meanwhile, research efforts on cell classification have also been evolving from fuzzy logic techniques [6], support vector machines (SVMs) [7] to cellular neural networks [8]. However, these works were mostly developed with a limited amount of data, and/or focused on images taken from specified instruments, which leave their generalization capability insufficiently justified.

Motivated by recent neuroscience findings [9, 10], unsupervised learning and deep learning techniques [11–13] have gained momentum during the past decade for object representation and recognition (e.g., face representation and recognition) [14–17]. And their applications in various biomedical tasks [18–22] have demonstrated success with the potential to provide a new avenue to data-intensive clinical studies, among which, the leukocytes classification accuracy has been significantly improved due to the employment of deep neural networks (DNNs) [23]. However, systems of such kind typically only provide the end-to-end (i.e., from data to classification) solution, which leaves the characterization of white blood cells inaccessible, and thus impede the construction of leukocytes profile for many potential needs, including profile interpretation, profile optimization, profile differentiation among cell types as well as profile association with other meaningful endpoints.

## Results

We applied unsupervised feature learning for the automatic acquisition of intrinsic signatures directly from raw data (microscopic images) that facilitates the efficient and effective characterization and classification of leukocytes, skipping conventional steps such as feature engineering (i.e., manually design and optimize features) while providing meaningful cell profile that is typically inaccessible to many existing deep learning systems. We show that a well-designed unsupervised learning system is capable of automatic and efficient extraction of signatures (i.e., patterns encode both color and texture information) directly from microscopic images, which provides the foundation for both meaningful representation as well as effective classification of white blood cells.

The general principle of our leukocytes characterization and classification system (referred to as WBC-Profiler) is illustrated in Fig. 1. WBC-Profiler is a hierarchical unsupervised feature learning framework with feed-backward reconstruction and feed-forward feature inference. Unlike many unsupervised feature learning algorithms [24–27], WBC-Profiler involves only element-wise nonlinearity and matrix multiplication. Therefore, it provides an highly efficient and effective solution for Leukocytes characterization and classification.

**Figure 1:**
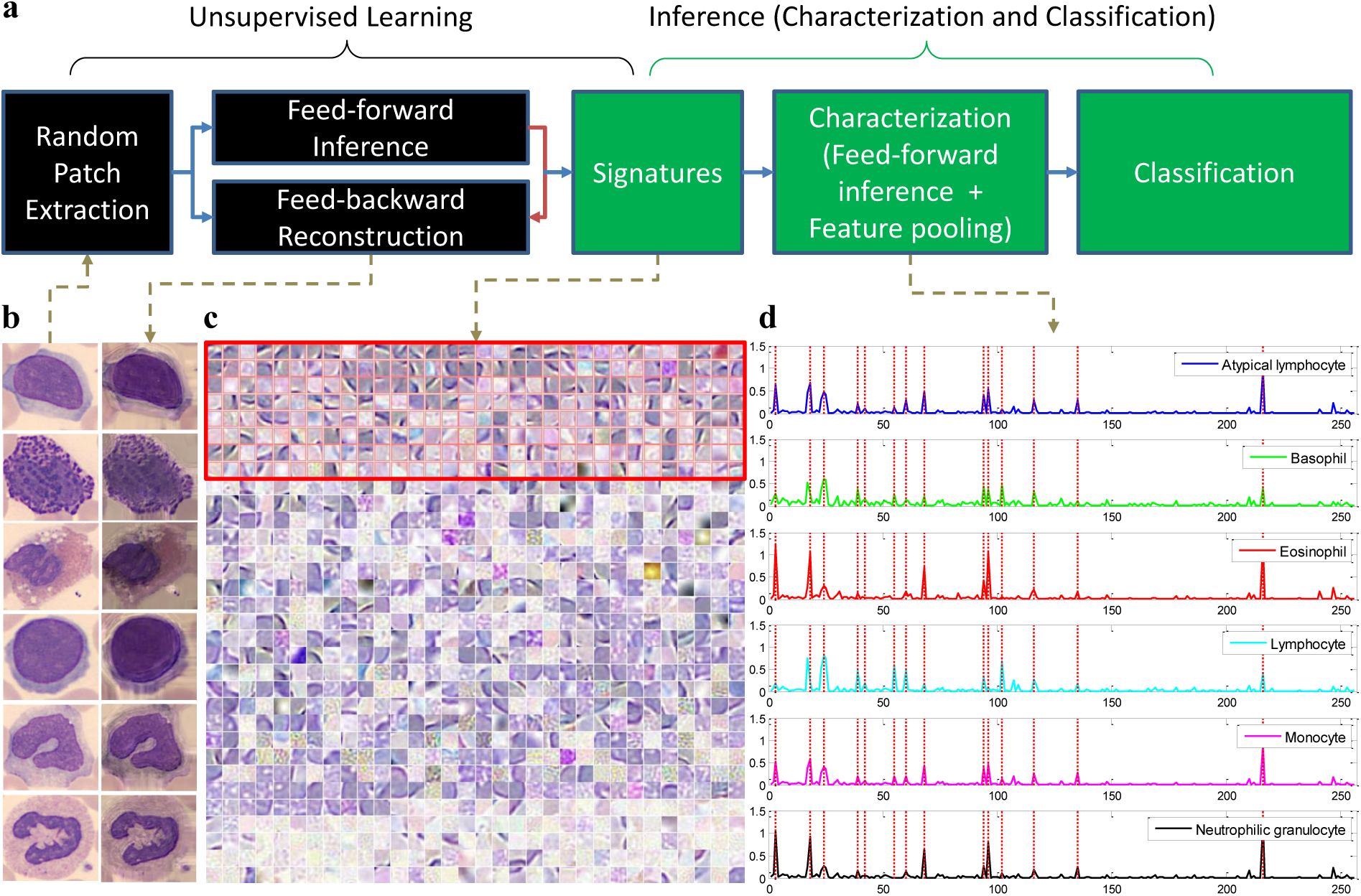
The concept of training and inference with WBC-Profiler for leukocytes characterization and classification. **a**, basic structure of WBC-Profiler, where the *random patch extraction unit* selects a set of vectorized image patches randomly from input cell images; the *feed-backward reconstruction unit* reconstructs the original input signal (e.g., image patch) from a set of signatures learned from the input data; the *feed-forward inference unit* predicts the reconstruction coefficient (i.e., sparse code to be used as the feature representation for each individual image patch) for input data reconstruction; the *characterization unit* calculates the sparse codes for all image patches of an target cell image based on both the signatures and inference function derived from unsupervised learning, and summarize them into a single profile as the representation of the target cell image; and finally, the *classification unit* labels each cell image with different cell types. **b**, Examples of microscopic images of different types of white blood cells and the corresponding reconstruction results from the derived signatures. From top row to bottom row: Atypical lymphocytes, Basophils, Eosinophils, Lymphocytes, Monocytes and Neutrophilic granulocytes. **c**, Signatures (1024 in total) automatically learned from our cell image dataset by WBC-Profiler, where the top 256 signatures (ranked by random-forest based on their contribution to leukocytes classification) were highlighted within red bounding box. **d**, Mean profile of the top 256 signatures per cell type, where it is represented as the class-average contribution of each signature for the reconstruction of cell images in the same category.

To evaluate the capability of WBC-Profiler on leukocytes characterization, we visualized the signatures acquired by WBC-Profiler from our leukocytes image database in Fig. 1c. It contains 1024 elements, and captures information from both color and texture domains, which cannot be easily achieved manually. With the derived signatures, WBC-Profiler extracts patch-level features for each image patch through a feed-forward fashion, where an image patch refers to a sub-image with fixed size (i.e., 20 pixel by 20 pixel in our study) cropped from the original microscopic image, and the patch-level feature of a single image patch refers to the sparse code (coefficients) for the reconstruction of this specific image patch with derived signatures. The final profile (image-level representation) for the entire microscopic image is constructed with specific pooling operation over all patch-level features from the microscopic image. The selection of pooling strategy is typically guided by the effectiveness of the corresponding profile during classification, and in our case, mean-pooling was selected, which constructs the profile as the average sparse code (i.e., average reconstruction coefficients) over all the image patches from the microscopic image. Heatmap of both selected signatures (see Fig. 2a) clearly indicates the differential expression of signatures across cell types, and provides an intuitive way to examine/evaluate the contribution of each signature for leukocyte image construction/composition. Furthermore, feature embedding of high-dimensional profile into 2-Dimensional (2D) space leads to cell-type-specific clusters (as illustrated in Fig. 2b-c), which demonstrates the effectiveness of WBC-Profiler in leukocytes characterization, especially for the task of leukocytes classification.

**Figure 2:**
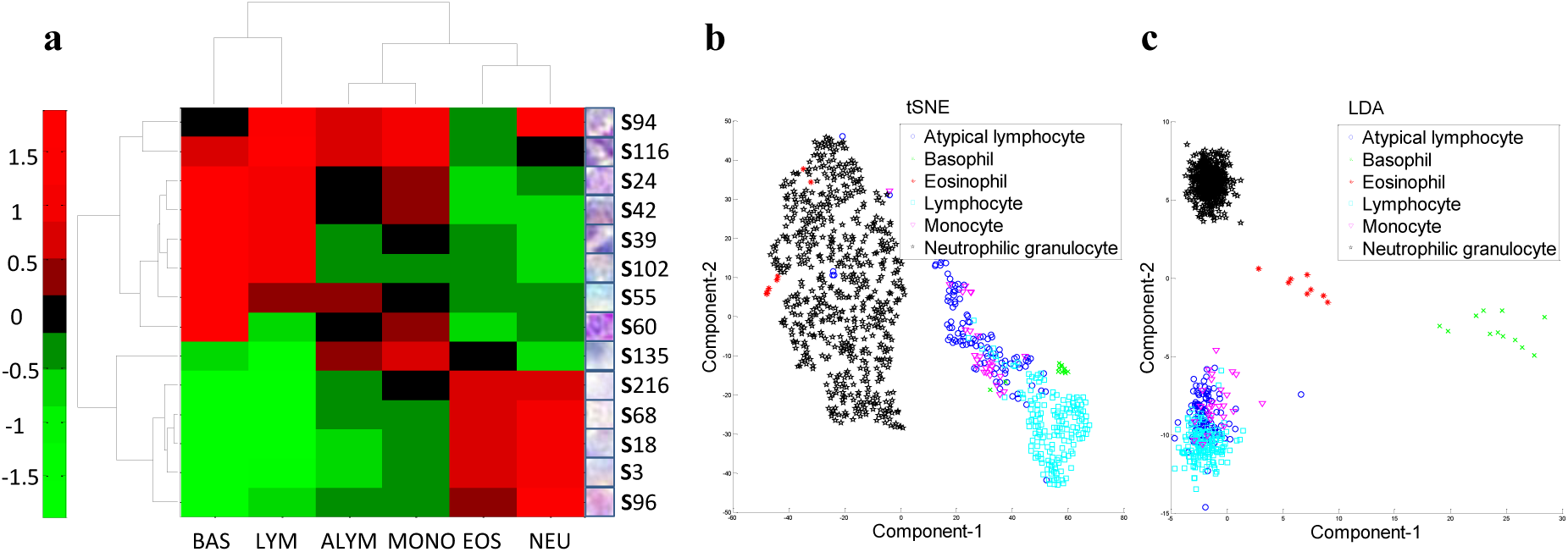
WBC-Profiler provides effective leukocytes characterization. **a**, Heatmap based on signatures selected from the mean profile (see Fig 1(d)), where zero-mean normalization is applied for better visualization. **b**, Feature embedding based on the top 256 signatures demonstrates the effectiveness of WBC-Profiler in unsupervised signature learning for leukocytes characterization, where t-Distributed Stochastic Neighbor Embedding (tSNE, unsupervised) reveals highly separable clusters. **c**, Separability of each clusters can be further improved through linear discriminant analysis (LDA, supervised).

To evaluate the capability of WBC-Profiler on leukocytes classification, we have compared it with one of the state-of-the-art techniques in the field of leukocytes classification (we refer it to as DeepVote, which combines the classification results from different deep neural networks to vote for the final decision) [23] and with one of the most successful deep learning techniques in the field of object detection and recognition, i.e., Faster R-CNN [28] with both vgg16 and res101 as the network architectures. During evaluation, half-half cross-validation was employed with 10 iterations, where, at each iteration, we randomly selected 50% of the data per cell type for training, and used the rest for testing, and the performance in terms of average F1-measure and confusion matrix was illustrated in Figure 3, which demonstrates the effectiveness of WBC-Profiler for leukocytes classification.

**Figure 3:**
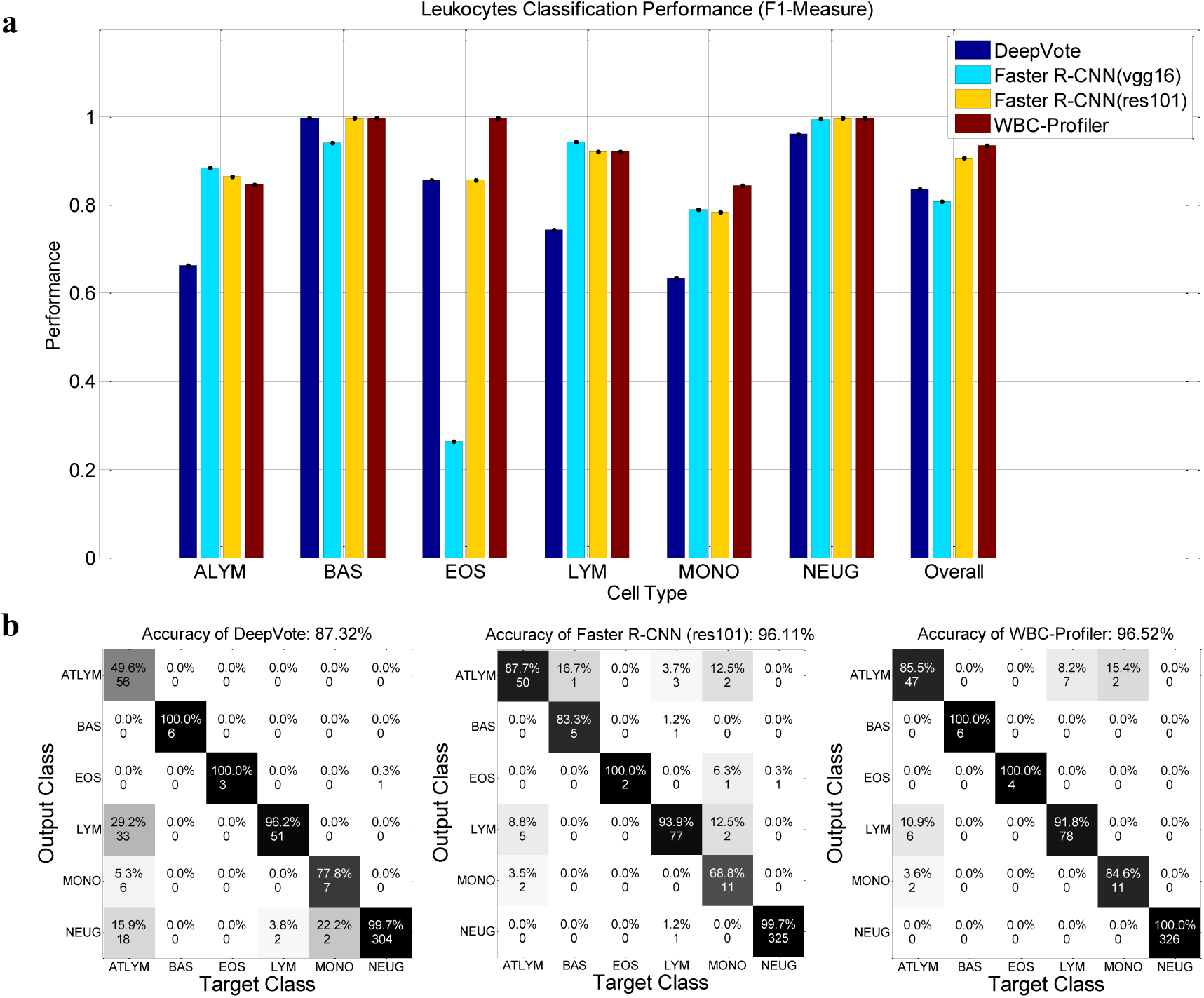
WBC-Profiler enables accurate leukocytes classification. **a**, Comparison of leukocytes classification performance in terms of F1-Measure for both individual class as well as overall class-average. **b**, Confusion matrix from DeepVote, Faster R-CNN (res101) and Stacked PSD for leukocytes classification with absolute cell image numbers (mean and rounded across all iterations) and normalized precision. Abbreviation: ATLYM - Atypical lymphocyte; BAS-Basophil; EOS - Eosinophil; LYM - Lymphocyte; MONO - Monocyte; NEUG - Neutrophilic granulocyte.

## Conclusions

In summary, we developed WBC-Profiler, an unsupervised feature learning system for efficient and effective leukocytes characterization and classification. As demonstrated through a well-curated dataset, complex signatures, capturing both color and texture information, can be automatically and directly learned from the microscopic images. The designed architecture and the feed-forward nature of feature inference ensures WBC-Profiler’s performance in precision, accuracy, and speed. Furthermore, WBC-Profiler decouples large scale complex leukocytes characterization from limitations in both human-based feature engineering/optimization and DNNs-based end-to-end solution, and therefore could further allow the extraction of highly multiplexed intrinsic properties and information from large scale leukocytes dataset towards meaningful endpoints with scientific and/or clinical impact.

## Acknowledgements

This work was supported by the Medical Key Science and Technology Development Projects of Nanjing (ZKX18016), the Medical Science and Technology Development Projects of Nanjing (YKK18167), and Synihealth Research.

## Author contributions

X.M. and H.C. wrote the software, performed the experiments, and analyzed the data. H.C., H.S., H.Y., and C.W conceived and supervised the study. H.C., H.S., and H.Y. wrote the manuscript. Y.X., X.X., R.X., Z.W., Z.L., Y.L, and X.Z created the leukocytes image database.

## Methods

### Leukocytes Image Database

A new dataset containing approximately 1000 microscopic images of 6 types of white blood cells has been collected at the Affiliated Drum Tower Hospital of Nanjing University Medical School to evaluate WBC-Profiler with detailed protocol listed as follows,

1. Sample selection. Abnormal blood test samples were selected by clinical experts for our study, where all samples were under daily intra-lab quality control as well as inter-lab quality control by the Clinical Lab Center, Jiangsu Province as well as MOPH (Ministry of Public Health) Clinical Lab Center.
2. Staining. Selected peripheral blood samples were filmed and stained with Wright-Gimsa through Sysmex SP 1000i.
3. Scanning. After staining, digital scanning was performed with OLYMPUS CX31RTSF at 1000X.
4. Labeling. The cell type labeling process were carried out by two clinical experts with more than 10 years of experience in the field.
5. Quality control. To ensure the data quality, only images meeting following criteria are selected:
  a. technical correctness. No artifacts were introduced during staining and scanning process.
  b. label consistency. The cell types of the target image independently labeled by the above two clinical experts were consistent with each other.
6. De-identification. To protect the privacy of patients, all images were de-identified by the clinical experts before any further quantitative study by WBC-Profiler.

### WBC-Profiler architecture

WBC-Profiler is composed of unsupervised signature learning module, inference module and visualization module to realize efficient and effective leukocytes characterization and classification, as well as convenient interpretation of derived results.

*The unsupervised signature learning module* of WBC-Profiler is build upon the Stacked Predictive Sparse Decomposition (Stacked PSD) technique [29] for the construction of hierarchical unsupervised learning framework, which is suggested to be able to capture higher-level dependencies of input variables, thereby improving the ability of the system to capture underlying regularities in the data. Unlike many unsupervised feature learning algorithms [24–27], the feed-forward feature inference of PSD is very efficient, as it involves only element-wise nonlinearity and matrix multiplication. Therefore, it provides an highly efficient and effective solution for Leukocytes characterization. In our study, we used only one PSD layer with 1024 kernels (signatures) at a fixed size of 20-by-20 pixels. The training process was set to be 100 iterations, and it converged with around 20 iterations.

*The inference module* of WBC-Profiler consists of characterization and classification units, where the former extracts patch-level features with a sliding window from the leukocytes image and construct the image-level profile through specific pooling operation, and the latter utilizes the pre-built profiles for leukocytes classification. In our study, we extracted patch-level features with a sliding window at the step size fixed to be 5 pixels, and adopted mean-pooling strategy among different popular pooling strategies through the evaluation on classification performance via support vector machine (SVM) classifier.

*The visualization module* of WBC-Profiler provides intuitive and convenient means for data visualization and interpretation, including signature visualization (as illustrated in Fig.1c), profile visualization (as illustrated in Fig.1d), heatmap (as illustrated in Fig. 2a) and feature embedding (as illustrated in Fig.2b-c).

### Code availability

Matlab scripts for WBC-Profiler, including unsupervised learning module, inference module and visualization module, are available as Supplementary Software, and further updates will be make available at http://bmihub.org/project/wbc-profiler.

### Data availability

The data that support the findings of this study are available from the corresponding author upon request. Example data are available in the Supplementary Data and Supplementary Software packages.

